# ATSASfold: a web server integrating AlphaFold models with ATSAS-based SAXS analysis

**DOI:** 10.64898/2026.11.18.767478

**Authors:** T. Tropin, A. Raasakka, C. Blanchet

## Abstract

Small-angle X-ray scattering (SAXS) is routinely used for the structural characterization of protein monomers in solution, and computational prediction methods such as AlphaFold now provide accurate atomic models for most proteins. When a predicted model and the experimental scattering curve disagree, several refinement strategies exist, but selecting the appropriate one remains difficult, particularly for users without SAXS modeling expertise. ATSASfold is a web server developed to help analyse the results of SAXS measurements on protein monomers and select appropriate refinement strategies. From a provided experimental SAXS curve, ATSASfold performs primary data analysis, gives a general assessment of macromolecules shape, size, possible aggregation state and data quality. An AlphaFold-generated model is fitted against the SAXS curve and the agreement with the experiment is evaluated. In cases where the fit shows significant discrepancies, a machine-learning classifier trained on SASBDB depositions assigns the case to one of four categories, suggesting the most relevant ATSAS-based strategy for model refinement: fixed model fitting, rigid-body modelling, normal-mode analysis or ensemble optimization. When needed, a graphical interface allows the user to adjust domains or flexible regions of the protein before submitting the structure to ATSAS online for longer fitting jobs. After the fitting is completed, ATSASfold provides a summary of the results. The workflow was tested on examples from the SASBDB open web-databank. These use cases demonstrate smooth operation of the web server and show how ATSASfold can simplify the transition from SAXS data collection to structure-based interpretation.

**Synopsis:** ATSASfold is web-server that integrates SAXS data analysis with evaluation of AlphaFold-predicted protein monomer structures and uses machine learning to recommend appropriate ATSAS-based refinement strategies when the models disagree with experimental data.

## 1. Introduction

Small-angle X-ray scattering (SAXS) is widely used for the structural characterization of proteins, other biological macromolecules, and their complexes in solution. SAXS provides reliable data on global structural properties, such as particle size and shape, molecular mass, and oligomeric state. Furthermore, SAXS allows the validation of structural models against experimental data. Yet, being a low-resolution method, reliable structural interpretation requires careful sample preparation, assessment of data quality, and data analysis [1].

The typical interpretation of SAXS data consists of several steps. It starts with primary data analysis – determination of basic structural parameters (e.g. gyration radius *R_g_*, maximum particle dimension *D_max_*) and evaluation of the quality of scattering data. Next, when an atomic model of the protein is available, its calculated SAXS profile can be compared with the experimental curve. In case of disagreement, different modelling approaches can be used. The choice of method depends on the origin of the discrepancy and on the studied macromolecule. The list includes rigid-body modelling of domains or parts of the complex, account of conformational changes for refinement, and treatment of flexible proteins as ensembles [2,3]. The ATSAS software suite provides established methods that cover all the stages of the described analysis, including primary data processing and structure-based modelling [4,5].

The recent progress in machine-learning-based structure prediction, marked by the introduction of AlphaFold 2 [6] and later AlphaFold 3 [7], has already significantly changed this workflow. High-quality predicted models are now available [8] and can provide an immediate starting point for interpretation of SAXS experiments. However, AF models still need to be validated against solution SAXS data. In particular, relative orientations of domains, flexible linkers, disordered regions and conformational heterogeneity in general may lead to significant differences between AF-predicted models and SAXS data. SAXS is therefore a useful experimental method for validating predicted structures and revealing discrepancies that require further modelling.

Several programs allow calculation and fitting of SAXS profiles from atomic structures, including CRYSOL [9], FoXS [10] and Pepsi-SAXS [11], while approaches such as MultiFoXS [12] and the recently introduced SAXS-A-FOLD [13] extend this analysis to flexible or ensemble models. AFflecto [14] provides another approach, generating conformational ensembles from AlphaFold models by sampling of flexible and intrinsically disordered regions, which can subsequently be evaluated against SAXS data. Another recent advancement is the BilboMD web-server [15], where users can follow a workflow to generate conformations that fit their data. Still, when a predicted structure does not fit the experimental data, selection of an appropriate modelling strategy remains a non-trivial step, particularly for users who are inexperienced with SAXS analysis.

Here, we present ATSASfold, a web server designed to guide this transition from an experimental SAXS curve and an AlphaFold model to an appropriate structure-refinement strategy for protein monomers. ATSASfold combines primary SAXS analysis with evaluation of the initial model-to-data agreement. In cases of significant disagreement, parameters obtained from the SAXS data and structural model are analysed using a machine-learning classifier to suggest an appropriate ATSAS-based strategy, including rigid-body modelling, normal-mode analysis or ensemble generation. The server further assists the user in defining structural regions required for subsequent calculations and provides access to longer ATSAS modelling jobs on the dedicated ATSAS online server. The workflow is demonstrated using experimental data and models deposited in the Small Angle Scattering Biological Data Bank (SASBDB), a curated repository for biological small-angle scattering data and structural models [16,17].

## 2. SAXS investigation of biomacromolecules

In this section a brief summary of the physics behind a typical BioSAXS measurement is given. The general experimental setup and parameters are covered, together with a recap of structural information obtained and the refinement strategies used to fit the SAXS curve.

### 2.1. General information on BioSAXS

In a SAXS experiment, a monochromatic focused X-ray beam is scattered by the randomly oriented macromolecules in solution, with the scattered intensity registered at the 2D detector positioned at a several meter distance behind the sample. The radially averaged intensity *I* is a function of the momentum transfer *q*:

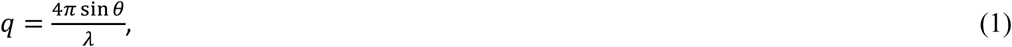

where λ is the wavelength of the incident X-ray beam and *θ* is half of the angle between the incident and scattered beams. After subtracting the scattering by the buffer solution, one obtains the *I*(*q*) function for all the further analysis. It is proportional to the orientation-averaged scattering of a single molecule in the solution:

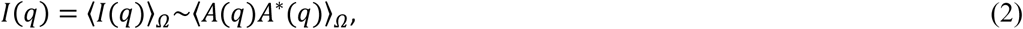

where *A*(*q*) is a Fourier transform of the scattering length density (SLD) *ρ*(*r*), that is proportional to electron density of the atoms.

A primary analysis of the *I*(*q*) curve allows extracting several structural parameters of the macromolecule. This usually comprises doing a Guinier analysis to extract *R_g_* and the forward scattering intensity *I*(0), calculating the pair distance distribution function (PDDF) to assess particle shape, and the Kratky plot analysis. From Guinier analysis one determines *R_g_* and *I*(0) via a low *q* approximation to *I*(*q*) (the Guinier approximation):

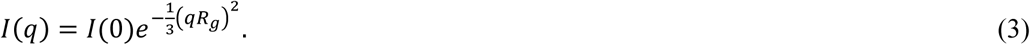

The *R_g_* and *I*(0) values can be conveniently extracted via Guinier plot, *ln I*(*q*) vs. *q*^2^ in the range of *qR_g_* < 1.3 (the general upper limit, that should be lower for non-globular particles). Attention must be paid to the linearity of the plot, which can be violated by mistakes in background subtraction, polydispersity, interparticle interactions, etc. The extracted *I*(0) value can be used to evaluate the molecular weight (*M_w_*) of the scattering particle.

The PDDF function is computed by a Fourier transform of the SAXS curve via:

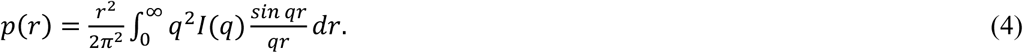

A perfectly spherical particle produces a bell-shaped *p*(*r*) with a maximum at *D_max_*/2. For anisometric particles this gets extended, while multidomain proteins may have several maxima in the *p*(*r*). Due to the limited range of experimental *q* and inevitable errors in *I*(*q*) values, computation of *p*(*r*) is not that straightforward, for details consult [18].

The folding state of proteins can also be evaluated from SAXS using the Kratky plot, *q*^2^*I*(*q*) vs. *q* [19]. The Kratky plot should decay at high *q* as ∼1/*q*^2^ for a folded protein, while for an ideal random coil it approaches a plateau. For a globular protein a peak at low *q* should be present, while for unfolded ones the peak is absent, and instead a continuous increase and a plateau will be observed. To compare between different samples, one should build the dimensionless Kratky plots, (*qR_g_*) *I*(*q*)/*I*(0) vs *qR_g_*. For a compact globular protein, the maximum of this reduced Kratky plot should be located at 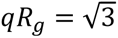 and 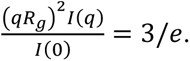

For an extended rigorous description of SAXS method applied to biomacromolecules we refer the interested reader to [20–22], and a more recent review [23].

### 2.2. Fitting SAXS curve

Given the atomic coordinates of a macromolecule, one can directly calculate the *I*(*q*) curve, either via the Debye equation, or by first computing the SLD *ρ*(*r*) and evaluating its Fourier transform, typically via multipole expansion. While for a protein in vacuum this results in a straightforward and direct task, for a solution one needs to account for excluded volume and the hydration shell scattering. The excess scattering of a particle in solution can therefore not be computed exactly from its atomic model alone. In practice, established approaches such as CRYSOL [9], FoXS [10,12], AUSAXS [24], absorb this uncertainty into two or three adjustable parameters, typically scaling the excluded volume and hydration shell density, which are optimized against the experimental curve.

If the fit is not satisfactory, given that other causes such as measurement errors, aggregation or interparticle interaction have been ruled out, this suggests that the proposed structure should be modified to correspond to experimental observations. To perform such modifications, the SAXS community has over the years developed a set of tools based on different approaches.

Several methods exist in ATSAS for *ab initio* shape determination, the main ones being DAMMIN/DAMMIF. The low-resolution model of the macromolecule is represented by densely packed beads in a constrained search volume and their arrangement is iteratively adjusted to fit the experimental data. With high resolution model, rigid-body modelling methods can be applied, typically SASREF, BUNCH or CORAL. They allow selecting the defined unchangeable parts of the molecule (rigid regions) and either just adjust their relative positions, or additionally model linkers/terminal structures in between. This significantly widens applications range of modeling using SAXS data.

Finally, for flexible and intrinsically disordered proteins (IDPs), the scattering does not correspond to a single shape, but to an average of multiple conformations that the protein samples in solution. In the ensemble optimization method (EOM), this average is approximated by a weighted average of a limited number of conformers: a large pool of random conformations is first generated with the help of a quasi-Ramachandran plot and a sub-ensemble whose averaged profile best fits the experimental curve is selected by a genetic optimization algorithm.

For detailed review of the mentioned hybrid methods, we refer the reader to [25], for ensemble-modeling to [26,27].

## 3. ATSASfold overview and user workflow

ATSASfold is designed as a web server providing a user-friendly GUI for users who are not necessarily proficient in SAXS investigations, analysis and modelling. The web-server was implemented using the Django framework. Substantial portions of the GUI and Django application code were implemented with use of OpenAI Codex under the direction of the authors.

The aim of the interface is to hide unnecessary complexity while still preserving access to the relevant tools and parameters. In addition, an “Advanced Mode” is available for more tuning at different stages of ATSASfold workflow. In this section we outline the general workflow of the website.

After registration and login, the user has an overview of the dashboard (Figure 1). The dashboard provides a simple list of user’s projects, sorted by recent progress. The ATSASfold **project** integrates the whole workflow corresponding to the analysis of a single SAXS measurement. Each user can run and manage multiple projects, conveniently switching between them via the dashboard.

**Figure 1.**
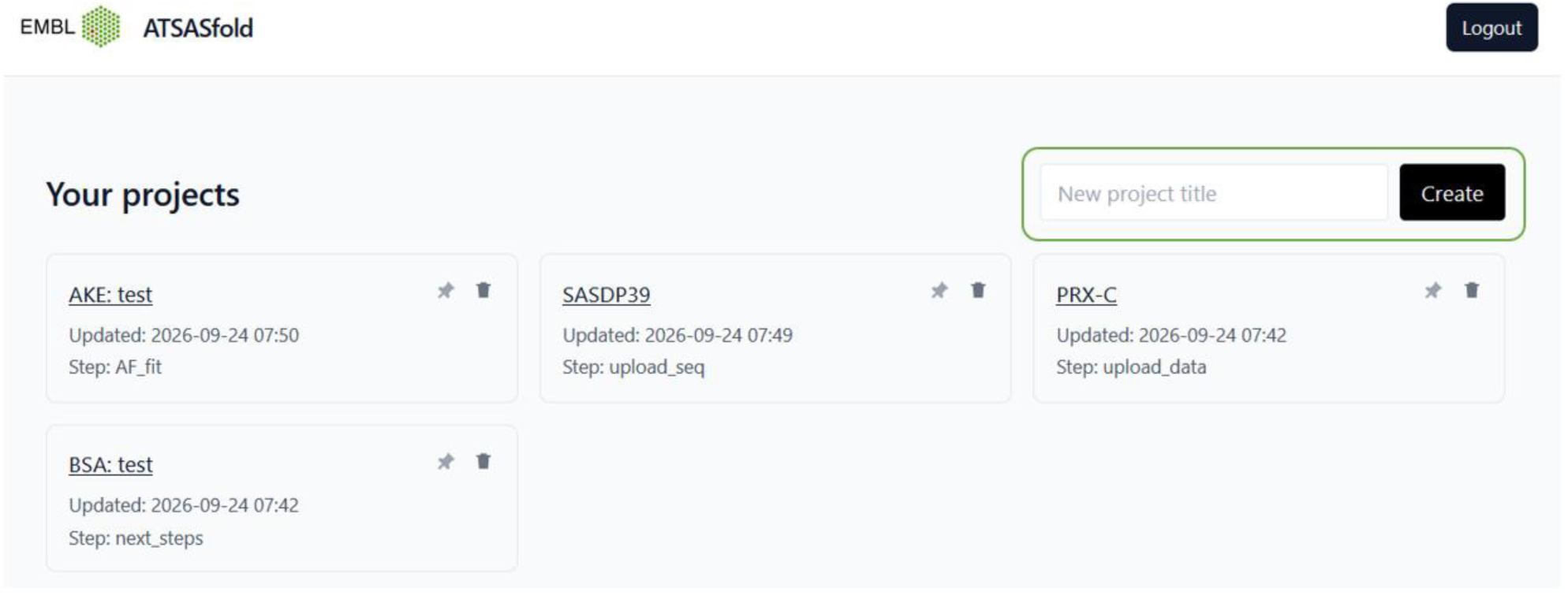
ATSASfold dashboard for user after logging in, with a list of projects (boxes), sorted by recent actions, and an input field for creating a new project.

Generally speaking, the project follows two main stages: (i) the SAXS data analysis, and (ii) the structure refinement. These parts are realised as two workspaces. While the movement through the project can be linear, the user can at any time return to the previous stages and adjust most parameters. Another option is to create a new project to start over with a new approach to data analysis. In the next sections the entire workflow is described in more detail, while here we give a brief overview of the interface layout and options.

After the project is created, the user is asked to upload an experimental SAXS curve of a protein in solution (the SAXS .dat file). The top-left panel displays the general steps of data analysis and fits, while the main panel shows the corresponding curves, distributions or 3D structures (Figure 2). If the step name is shown in green, then the main part of the analysis connected with it has been completed. Otherwise, the step name is shown in gray meaning that either ATSASfold is waiting for user action, or the analysis is still proceeding at the server. When some user action is required, the upload boxes are shown on the right panel together with short instructions. As analysis results are generated, the top-left panel is extended down with tabular information (Figure 2).

**Figure 2.**
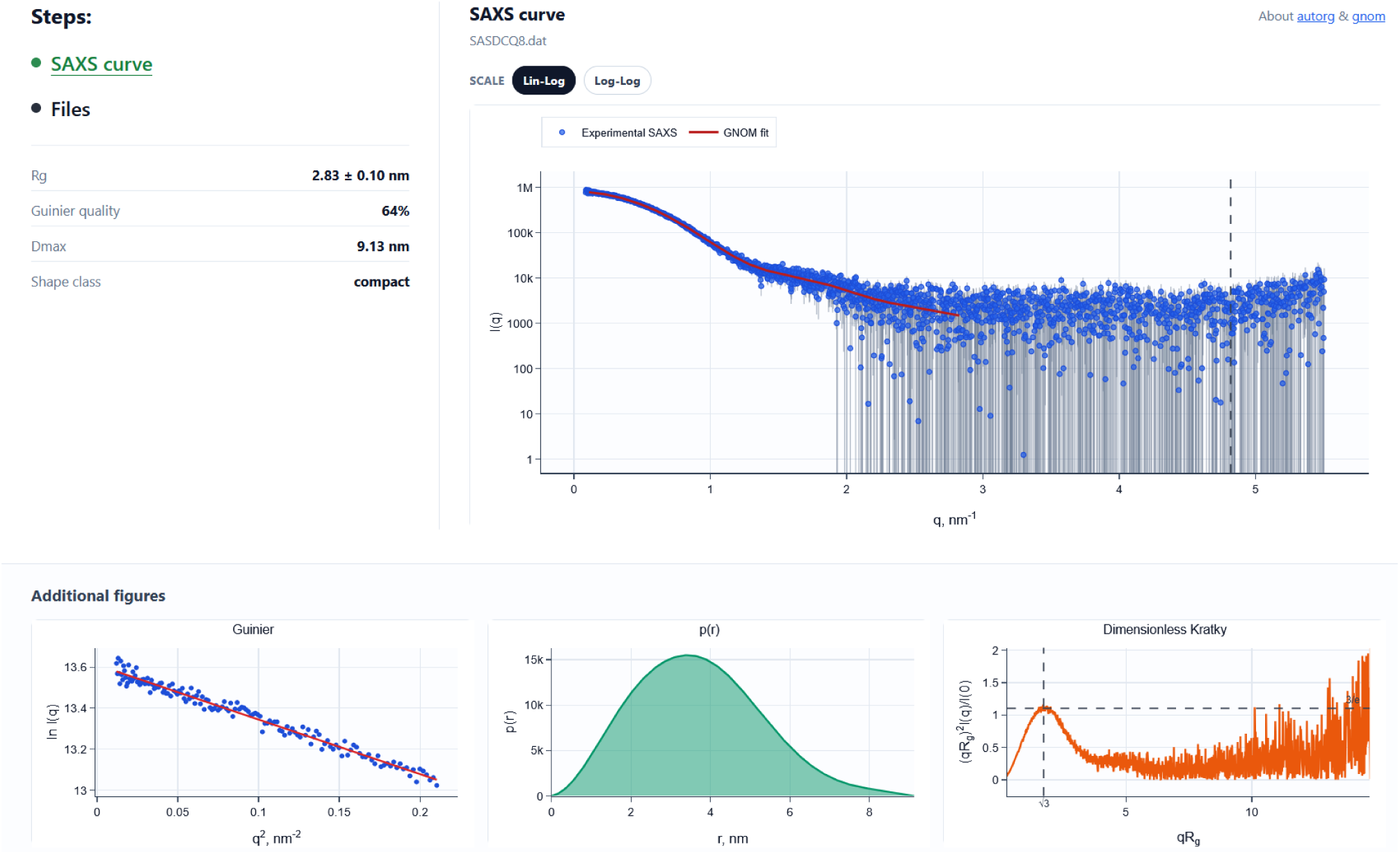
SAXS analysis interface. In the top-left corner consecutive steps of the workflow are presented. Other parts of the page present analysis results or query user to upload data files.

The additional bottom panel displays supporting graphical information such as the Guinier fit, the PDDF and the dimensionless Kratky plot. The individual steps of the SAXS data analysis are described in more detail in the following section.

The second workspace of ATSASfold guides the user through the structure refinement part. This workspace consists of two tabs – the domain editor and the ATSAS runs (Figure 3). The left main panel displays the 3D model and respective interface for domain auto-estimation and modification (domain editor). It can also be toggled to display the status/results of one of several possible ATSAS online runs. In the domain editor mode, the user can either run automated tools for domains generation or manually adjust the domains boundaries, linkers and termini. This is done either by dragging the boundaries on the pLDDT plot with the mouse, or manually changing numbers in the bottom panel, or adding/deleting new domains via action buttons. The manually changed domains configuration must be saved (by pressing the “Save Domains” button) before submitting to ATSAS online.

**Figure 3.**
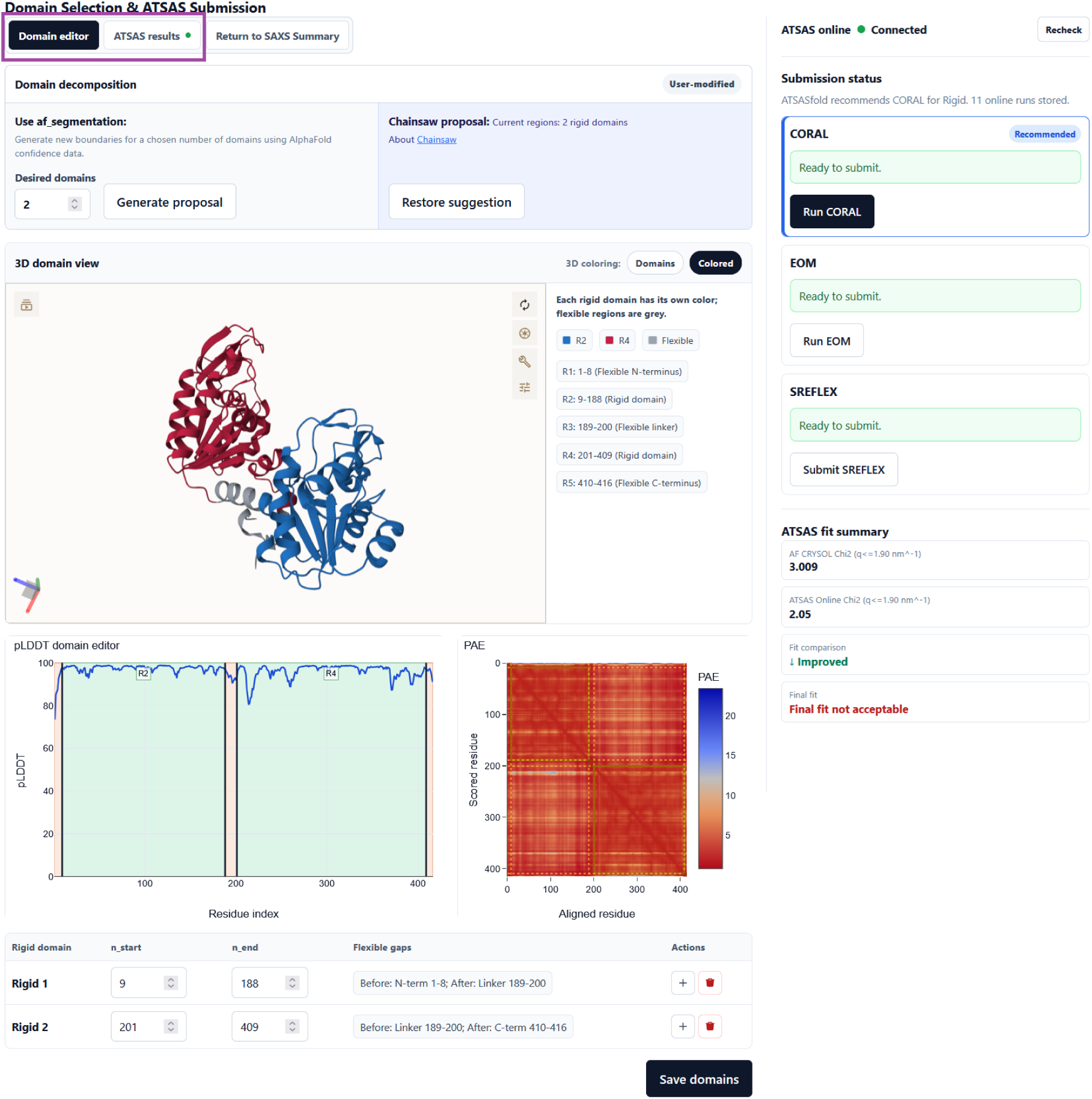
Domain editing & ATSAS run submission workspace. Two tabs at the top (outlined in purple) allow switching between selecting domains and checking the status of ATSAS online projects. The domain selection is guided by the displayed 3D structure, PAE and pLDDT scores. On the right-hand side, a list of available methods is given with the ATSASfold recommended one highlighted by a blue box.

The right panel of the domain editor is an interface for submitting the project to ATSAS online. In the “ATSAS results” tab, on the other hand, this panel allows opening the ATSAS online project in a separate browser tab (user must be logged in to the ATSAS online website), or cloning a project to the domains editor. More importantly, when the calculations on the ATSAS online servers are finished, this panel displays the results. The initial 3D model is shown in comparison with the new model obtained after fitting. Below, the SAXS fit results are presented for the user. The right-hand side shows the ATSAS fit summary with a comparison of initial and new *χ*^2^ values. This panel also allows downloading a zip-file with the ATSAS online results.

Finally, when finishing work with the project, a user can download fits, parameter tables, and refined structures to his PC for further investigation.

## 4. Methods and implementation

With the ATSASfold workflow overviewed in the previous section, in this section we describe the steps of the SAXS data analysis in more detail.

### 4.1. Primary SAXS data analysis

After the experimental SAXS curve is uploaded, ATSASfold performs primary data analysis using methods from the ATSAS suite (autorg, datgnom, datclass, datmw). The ATSAS tool calls are performed via the pyatsas Python package [28]. A scheme of the primary analysis run is presented in Figure 4. The standard SAXS parameters such as *R_g_*, shape (class) and molecular weight are computed. First, autorg identifies the Guinier interval and estimates *R_g_*, *I*(0) and the Guinier fit quality. The first retained data point is taken from this interval, while shanum is used later to select the last point (via the informative *q*-range) and to estimate the number of Shannon channels [29]. datcrop is then used to generate the curve for subsequent calculations in ATSASfold. Both the Guinier interval and *D_max_* can be adjusted manually in advanced mode, after which the dependent analysis stages are recalculated.

**Figure 4.**
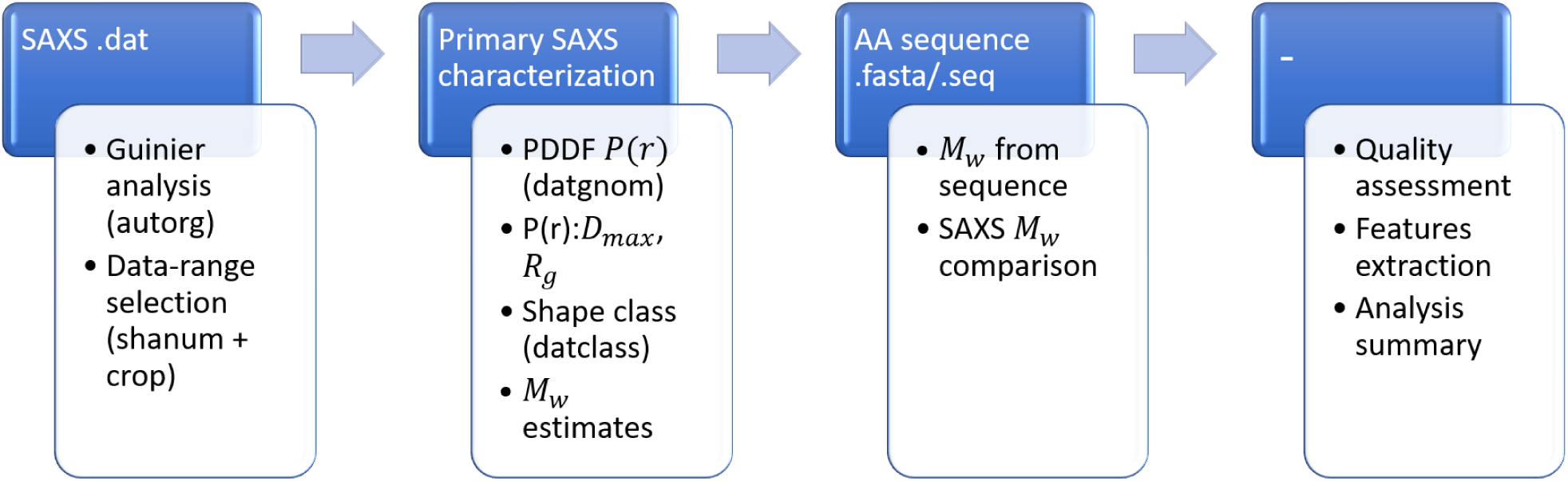
Scheme of ATSASfold initial SAXS data analysis. The blue boxes reflect input required from the user (last step – no input), the white boxes show the corresponding steps performed by ATSASfold.

The cropped curve is analysed using a few complementary methods: (i) datgnom performs the indirect Fourier transform (IFT) and provides *P*(*r*), *D_max_* and real-space estimates of *R_g_* and *I*(0); (ii) datclass assigns a general particle-shape class; (iii) datmw calculates molecular-weight estimates using Bayes, Porod, *Q_p_*, *V_c_*, MoW, and size-and-shape approaches [4,30]. Then, the experimental uncertainties are additionally evaluated against the datgnom fit, so that revised uncertainties can be suggested when the supplied errors are missing or inconsistent.

After a protein amino acid sequence is provided, the theoretical molecular weight is compared with the SAXS-derived Bayesian estimate (or with the Porod estimate when Bayesian is unavailable). Agreement within 20% is reported as consistent. A SAXS molecular weight 20–50% higher than the sequence value may indicate aggregation, whereas a difference above 50% is interpreted as possible oligomerization and used to estimate the number of subunits. A significantly lower SAXS *M_w_* estimate may indicate an incorrect sequence, truncation, or poor data quality.

The extraction of features for the ML analysis (see below) starts also during this stage. The analysis details, the model and list of features are reported in the Appendix A.

The obtained results provide the first step before any structure-based fitting is performed. This allows the user to assess SAXS data quality and gain initial structural information. A summary is displayed to the user: the tabular data together with Guinier, PDDF, Kratky plots. Warnings on data quality, aggregation or oligomerization are also displayed when needed.

### 4.2. AlphaFold model and initial SAXS fit

An AlphaFold model can be supplied as a structure file together with PAE and pLDDT scores, or as a complete prediction archive (.zip file). For authorized user, there is also a limited option of submitting the protein sequence to the configured AlphaFold run on the cluster. Alternatively, a local protein sequence similarity search can be performed using BLASTP [31]. It identifies candidate entries in the AlphaFold Protein Structure Database (AFDB) [8], after which the highest-ranked downloadable structure and PAE data are retrieved. This search identifies closely related candidates and does not require 100% sequence identity. Nevertheless, providing an AlphaFold structure via inference on the given protein sequence is preferred for subsequent work in ATSASfold.

The selected structure is fitted to the SAXS curve using CRYSOL. Fits are performed for several different *q_max_* values. Next, the default fit is evaluated with datcmp using CorMap, Chi-square, and Anderson–Darling tests [4,32]. The fit is considered acceptable when two of three tests satisfy the acceptance criteria. The calculated and experimental curves, CRYSOL *χ*^2^, test statistics, and associated p-values are displayed to the user.

If the AlphaFold model fits the SAXS data well, the workflow can stop with a final summary. If the discrepancy is significant, ATSASfold proceeds to the next steps.

### 4.3. Four-class ML classification

In order to proceed to structure refinement, ATSASfold first runs a ML-based classifier to assign the project to one of four practical categories. The classifier output is used to suggest whether the user should try rigid-body modelling, normal-mode analysis, or ensemble modelling. An XGBoost classifier [33] trained on the dataset composed of downloaded and processed SASBDB depositions is used (see Appendix for details).

The suggested categories are not intended as a final structural interpretation, but as a recommendation for the next modelling route. Additionally, to guide the user, the per-class probabilities are reported (Figure 5). The four categories are:

1. **Direct fit** – AlphaFold-generated protein structure fits the SAXS data sufficiently well. No further refinement is needed.
2. **Rigid-body** – rigid-body modelling is advised for improving SAXS fits.
3. **Normal-mode** – normal-mode analysis is advised for refining protein structure to match experimental SAXS data.
4. **Ensemble** – a single structure is not sufficient to fit SAXS data. An ensemble of structures should be generated.

**Figure 5.**
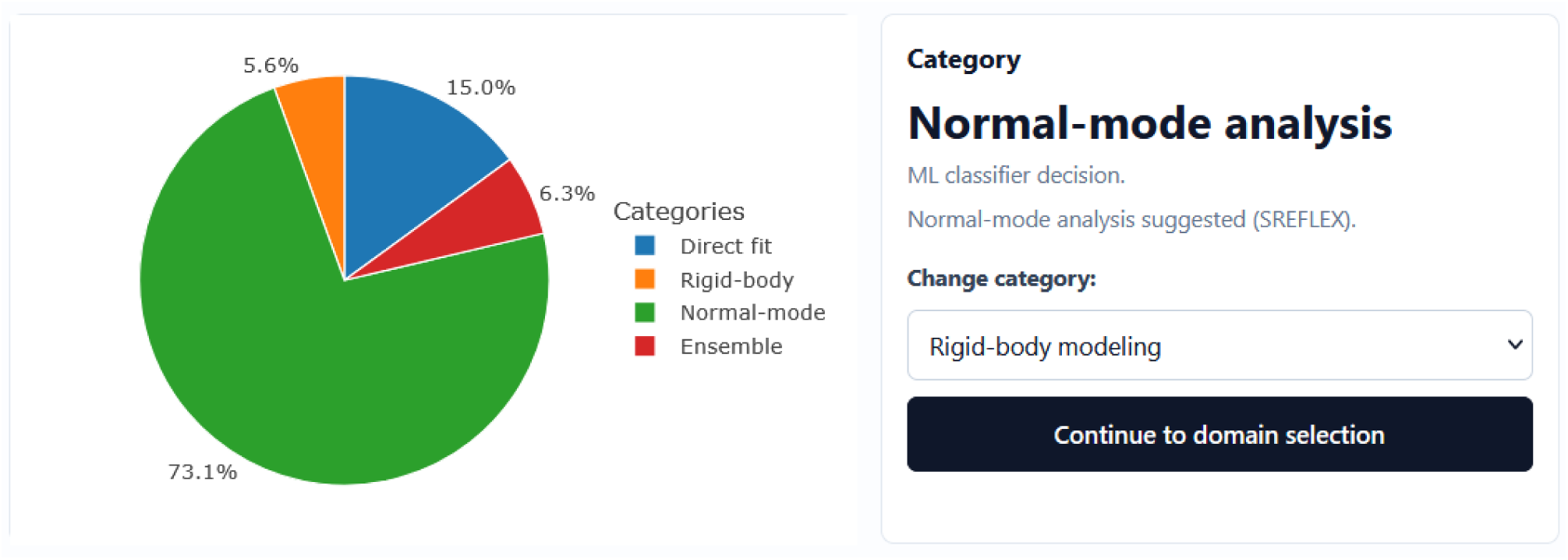
Left: pie-chart representing ML-classifier run results in terms of per-class probabilities. Right: option to override category choice.

The user has the option to override ML-chosen category and proceed with any of the available scenarios for further fitting.

### 4.4. Domain selection algorithm and GUI

For proteins composed of several domains, or a single domain with flexible termini, ATSASfold provides a graphical interface for domains selection. Behind the scenes, domain identification is performed by three different approaches: ATSASfold’s own tunable segmentation (af_segmentation), Chainsaw, and Merizo. Both Chainsaw and Merizo are supervised deep learning approaches to predict domain boundaries with high accuracy. Merizo [34] uses an encoder-decoder architecture with invariant point attention (IPA) to cluster residues into domains. It was evaluated using a test set comprising only multi-domain proteins. However, it is designed to handle both single- and multi-domain architectures. The other approach, Chainsaw [35], utilizes a fully convolutional neural network (2D CNN), trained on a balanced dataset of proteins (both single- and multidomain) ensuring versatility in domain boundary prediction. Additionally, unlike Merizo, Chainsaw explicitly reports the flexible linkers between domains.

The ATSASfold domain segmentation method, af_segmentation [36], relies on its own combined boundary score. The method has the advantage of searching for a user-defined number of domains and thus can be used as an additional advisory tool. The boundary score is mostly governed by the local PAE score and can be further adjusted via the low-pLDDT contribution.

This local PAE boundary score is calculated as:

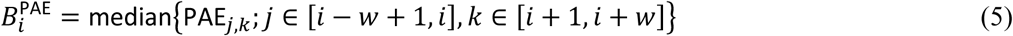

(also not expressed here is the clipping to stay in the (0, *L*) boundaries). Thus, this is a median PAE between residues in the left and right windows. The main governing parameter is *w* – the window size. Higher *B^PAE^* means that the two sides have an uncertain relative position, signaling that this is probably a boundary. To adjust for the low-PLDDT contribution the combined score includes the pLDDT score according to:

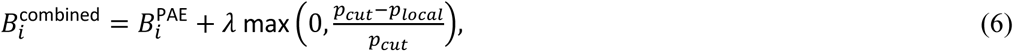

where *λ* weights the pLDDT input, and *p_cut_* defines pLDDT confidence threshold. Note that here *p_local_* is not just pLDDT *p_i_* itself, but the median pLDDT calculated over the *w_pLDDT_* window. For the balanced automatic mode of the ATSASfold segmentation the following parameter values are used: *p_cut_* = 60, *λ* = 8, *w* = 12, *w_pLDDT_* = 4.

The general workflow of domain segmentation is as follows. Chainsaw and Merizo predictions are generated and cached in advance (once atomic structure becomes available to ATSASfold). The Chainsaw prediction is taken by default if its confidence score is greater than or equal to 0.8. Otherwise, if Merizo score is greater or equal to 0.7, this suggestion is taken. For all other cases af_segmentation is used as a fallback.

The predicted domains are converted into rigid regions, while uncovered residues and short regions between domains are taken as flexible linkers. The user can review and manually edit the domain regions. Alternatively, the user can use af_segmentation to generate a new proposal for a chosen number of domains. The user can also manually modify, remove or add domains via the GUI, as described previously.

Overall, this step prepares the submission model for ATSAS tools that require domain ranges as input.

### 4.5. ATSAS Online submissions

Calculations involving ensemble averaging, rigid-body modelling or normal-mode analysis can be seamlessly delegated to run on the ATSAS Online server [37]. A prior registration is necessary: after the user registers in ATSASfold with the same email as in ATSAS online, submission of projects becomes available.

Once the domains have been selected, the user does not need to manually prepare all the input files. The ATSASfold backend automatically prepares the submission files together with the .pdb files for the corresponding rigid domains. It can query ATSAS online, create a project, submit the SAXS curve and other required files. Further, ATSASfold tracks the submitted job and retrieves the zipped results when the calculation is finished. Once finished, the user is presented with the results of the optimization run. The initial and refined SAXS fit and the corresponding *χ*^2^ values are displayed. For rigid body modelling the new 3D structure is compared with the initial one. For ensemble optimization, the selected conformations and ensembles (via their *R_g_* distribution) are also presented.

### 4.6. Summary

At the end of the workflow ATSASfold generates a summary containing the primary SAXS analysis, model-fitting results and suggested interpretation. The pdf report includes plots of experimental versus calculated scattering, residuals, fitted structural models and key numerical parameters. Output files are also downloadable as a zip package so that user can continue analysis manually, compile results for a publication, or upload the models to SASBDB together with the SAXS data.

## 5. Use-case examples and validation

The operation of ATSASfold has been tested with different SAXS datasets. Specifically, the quality of the primary analysis, the domain identification and editing, the submission of modeling jobs to ATSAS online and the returned results were tested. In most cases the data and models obtained from SASBDB entries were used. Selected examples are presented below, illustrating different pathways of ATSASfold operation. First one show an acceptable AlphaFold structure fit, second one a route that requires rigid-body analysis and discusses some controversial results, while the final example shows fitting SAXS data of an IDP with an ensemble of conformations.

### 5.1. AF model fit to SAXS data

As a straightforward example of fitting an AlphaFold model to experimental SAXS data, the SEC-SAXS measurements of bovine carbonic anhydrase II were used as input (SASBDB dataset SASDFP8 [38]). Carbonic anhydrase II is a compact globular protein with a molecular mass of ∼29 kDa. The SASBDB entry reports an experimental *M_w_* of 28 kDa, *R_g_*∼1.8 *nm*, *D_max_*∼5.1 *nm*.

The primary SAXS analysis performed by ATSASfold indicates a compact, folded particle, as evidenced by the Guinier, dimensionless Kratky and PDDF plots (Figure 6). The AF predicted a compact structure with high confidence over the structured part of the protein (mean pLDDT score per residue > 95). Direct calculation of the scattering profile from the AF model with CRYSOL results in good agreement with the experimental data (*χ*^2^∼1.03). Given this information, the ML classifier correctly and confidently suggests the “Direct fit” class and no further structure adjustments (Figure 7, SASDFP8 was excluded from the dataset for training the model for this demonstration). This example demonstrates the basic ATSASfold workflow for the case where a single, well-defined AF structure provides an adequate representation of the protein conformation in solution.

**Figure 6.**
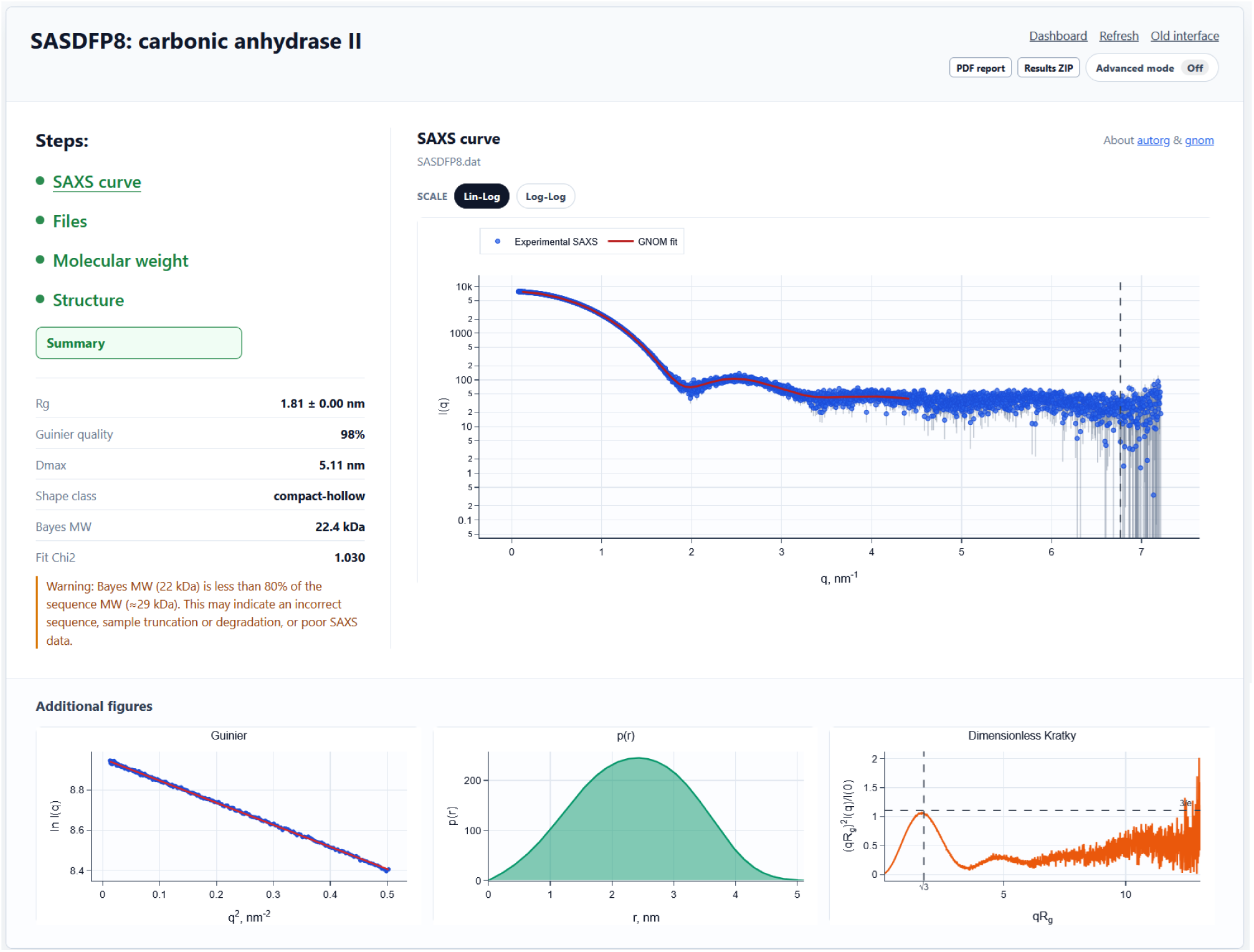
SAXS curve panel of ATSASfold for carbonic anhydrase 2 (SASDFP8 [38]).

**Figure 7.**
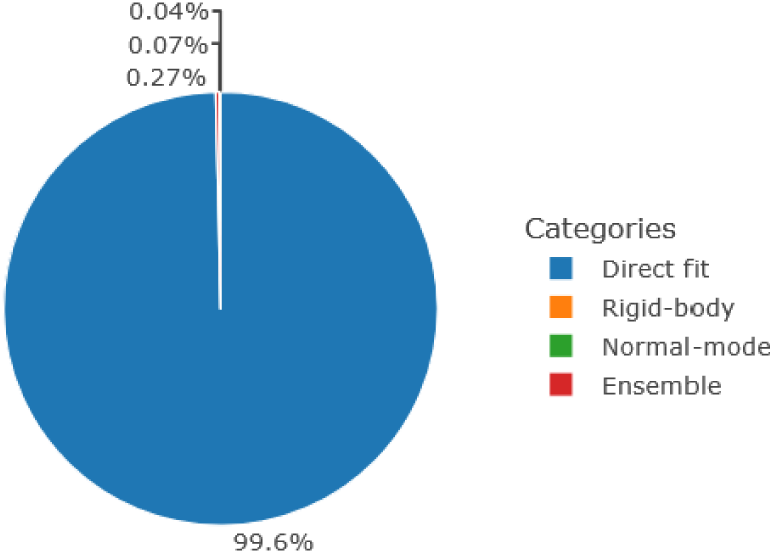
Summary confidence scores for SASDFP8, together with *χ*^2^∼1 suggest that no further adjustments required.

### 5.2. Structure refinement example

Arginyl-tRNA–protein transferase 1 (ATE1) from *Saccharomyces cerevisiae* (ScATE1) is chosen here to demonstrate the SAXS-guided refinement of an AlphaFold model using rigid body modelling. ATE1 catalyzes tRNA-dependent post-translational arginylation, including the addition of arginine to N-terminal residues in substrates of the Arg/N-degron pathway, thereby contributing to regulation of protein stability and other cellular processes. ScATE1 has a bilobed architecture comprising an N-terminal regulatory domain, a GCN5-related N-acetyltransferase (GNAT) fold and a C-terminal domain.

The SASBDB deposition SASDP39 [39] has an integrated SEC-SAXS curve for monomeric apo ScATE1, with *M_w_* = 59.9 *kDa* (experimentally estimated ∼62 kDa). The Guinier and PDDF-based analysis of SAXS data gives an *R_g_* of 2.8 nm (AF2-structure *R_g_*∼2.5 nm) and a *D_max_* of 8.3 nm.

The SASDP39 data were collected as part of the structural characterization of apo ScATE1 by X-ray crystallography, SEC-SAXS and cryo-EM [39]. Structural analysis places the catalytic region and the proposed tRNA-binding site around the GNAT fold and the interface between the two lobes. In the apo state, the N-terminal region is comparatively dynamic; this domain has also been implicated in binding a regulatory metal cluster. The AlphaFold prediction agrees closely with the crystallographic structure, but the SAXS data indicated that the relative arrangement of these regions in solution was not reproduced optimally (Figure 8, left). ScATE1 therefore provides an example in which an AlphaFold model is locally highly accurate while its inter-domain arrangement requires adjustment to describe the solution-state.

**Figure 8.**
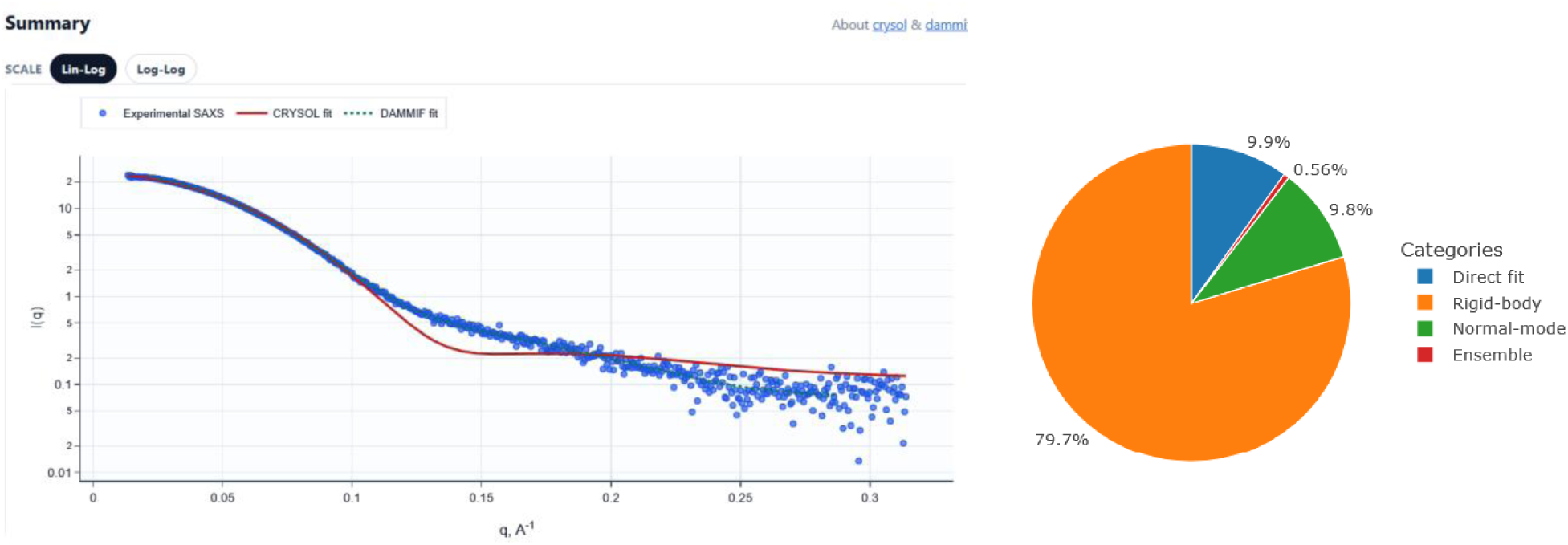
**Left** – experimental SAXS curve of ATE1 (blue circles; SASDP39 [39]) with CRYSOL fit of obtained AlphaFold structure (red line), as presented in ATSASfold interface. The calculated *χ*^2^ is 9.6. **Right** - ML classifier decision pie-chart strongly suggesting the rigid-body analysis method.

With an initial SAXS fit of *χ*^2^ = 9.6, the ATSASfold classifier’s suggestion is to use rigid-body modeling for the fit (Figure 8, right), with ∼10% probabilities for the Direct fit and Normal-mode classes. Interestingly, in the original study [39], normal-mode refinement with SREFLEX was used to relax the tailless AlphaFold2 model (residues 1-503), improving the fit 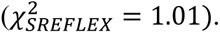 However, on the full experimental construct, SREFLEX cannot detach the folded-back C-terminal affinity tag from the protein surface. Instead, it attempts to achieve the required increase of *R_g_* by displacing the rigid normal-mode blocks and introducing backbone discontinuities; moreover, stalling at *χ*^2^∼2.2 in restrained refinement. Notably, inspection of the originally deposited SREFLEX model (fit 2 in SASDP39) reveals the same normal-mode partitioning artifact, including the tearing of a *β*-strand out of the catalytic GNAT *β*-sheet.

By contrast, following the ATSASfold classifier’s suggestion and using domain-aware boundaries (as determined by Chainsaw), a CORAL refinement results in a good fit (Figure 10 Right, *χ*^2^ = 1.44, *R_g_* = 2.78 nm) with no C_α_ clashes or domain distortions [40]. Structurally, this model preserves both the intact GNAT active site and the composite N/C-terminal β-sheet core while opening the inter-lobe cleft across the regulatory loops observed to be disordered in the apo *Sc*ATE1 crystal structure (Figure 9 Left, and pdb entry 7TIF). This also directly supports the allosteric cleft-opening mechanism proposed in [39].

**Figure 9.**
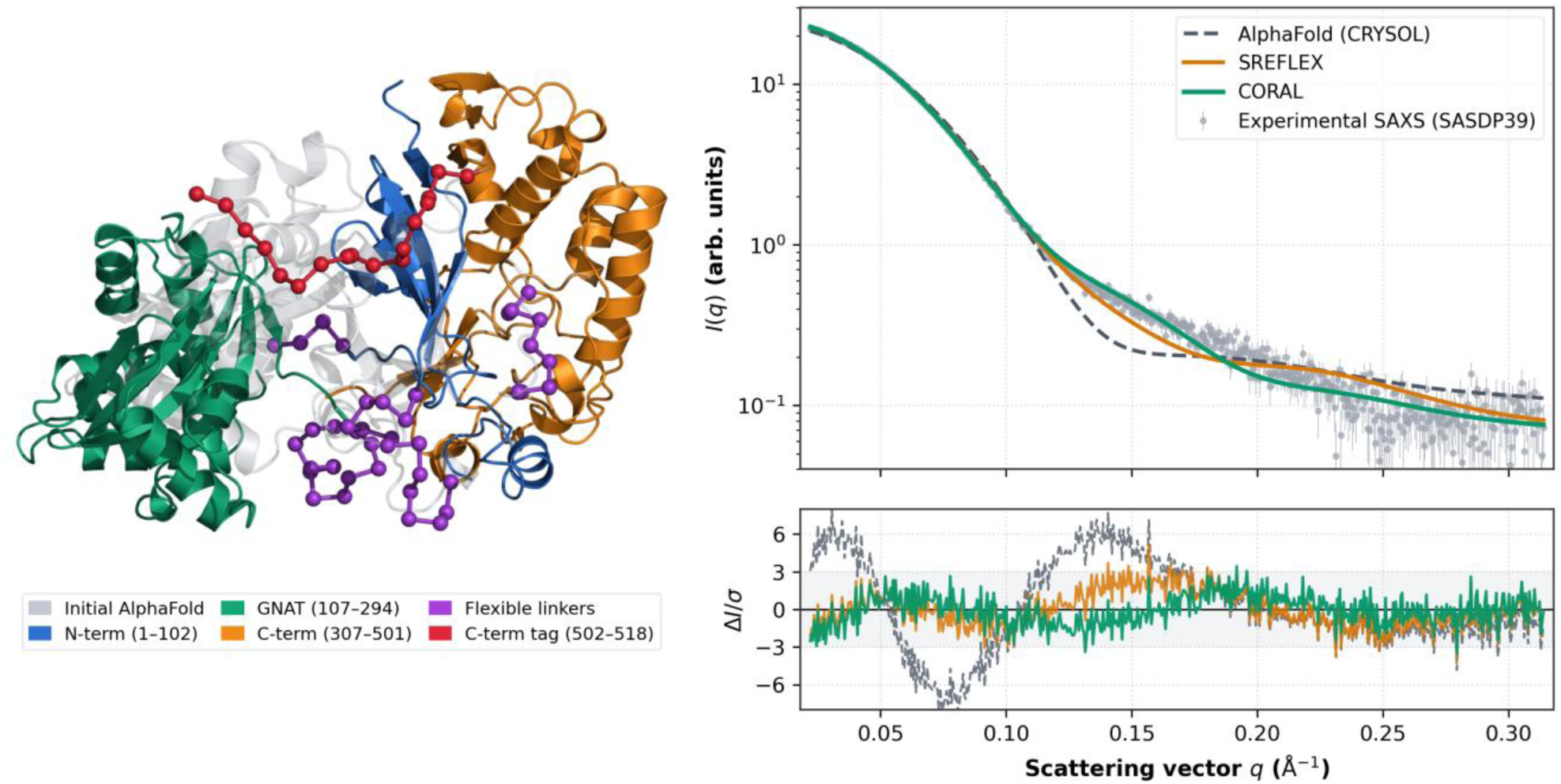
**Left** – initial AlphaFold structure (gray) and CORAL-refined domain-aware model (multicolor; flexible linkers and affinity tag shown as C_α_ spheres) of apo ScATE1. **Right** – SAXS data: experimental results (gray circles), fit of the AlphaFold structure (dashed gray line), SREFLEX (orange) and final CORAL model (green), with error-weighted residuals shown below.

**Figure 10.**
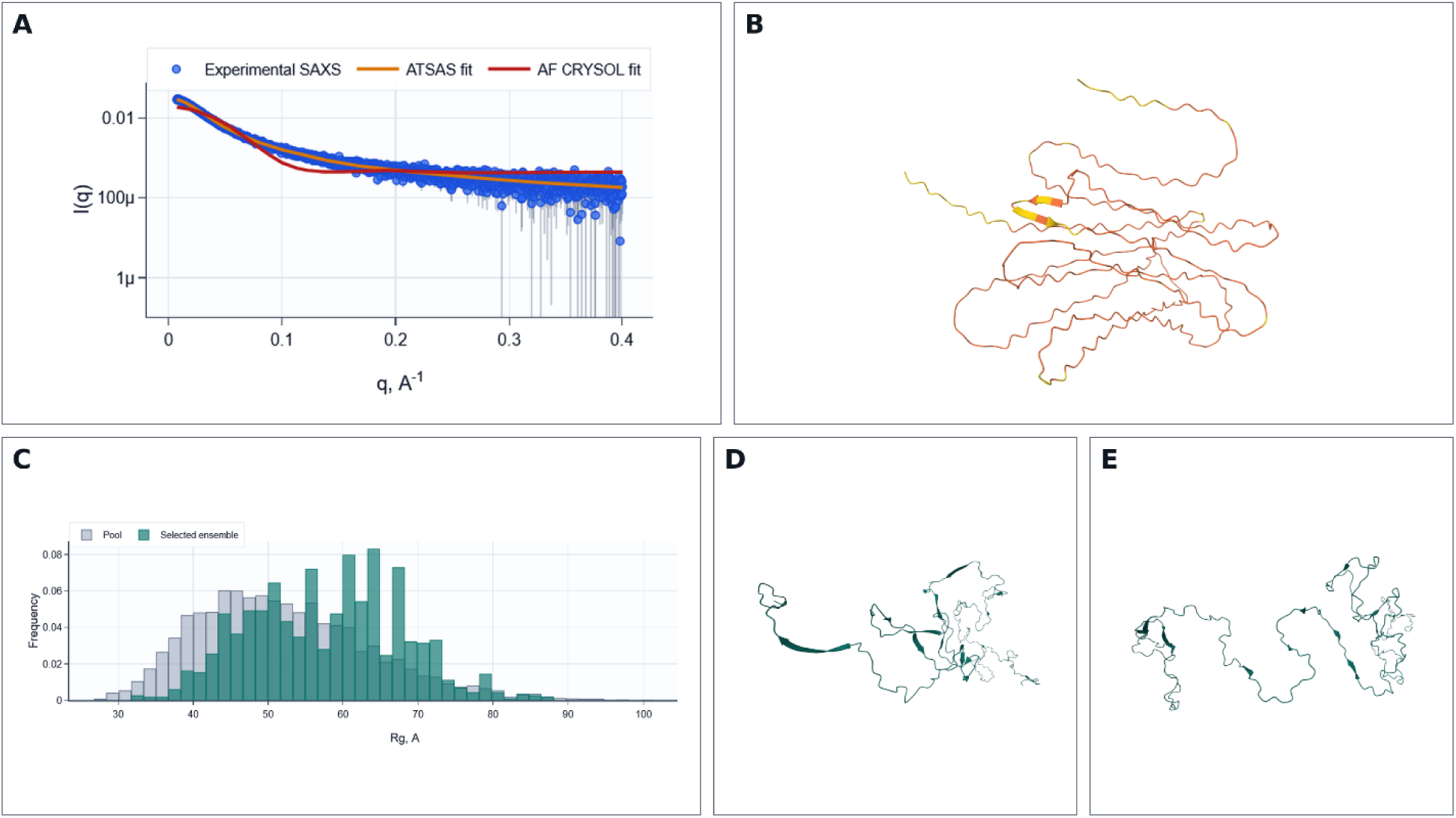
**A** - SAXS experimental data for PRX-C (SASBDB entry SASD2N7 [41]), together with the CRYSOL fit of AF-generated structure, and the results of an EOM run. **B** - AlphaFold-generated structure, with orange color corresponding to pLDDT score below 50. **C** - The *R_g_* distribution of ensembles initially generated by EOM (“pool”) and the representatives selected for SAXS fitting. **D, E** – selected structures from EOM fit pool having *R_g_* values of 5.6 nm (D) and 6.9 nm (E).

### 5.3. Ensemble modeling to fit SAXS curves

As an example of modelling of an intrinsically disordered protein (IDP) based on SAXS data, we consider the case of the C-terminal region of periaxin (PRX-C) (SASBDB entry SASD2N7 [41]), derived from the long isoform of periaxin (L-PRX). L-PRX is a long dimeric protein that is an abundant component of non-compact peripheral myelin, where it participates in molecular assemblies connecting the Schwann cell cytoskeleton with the extracellular basal lamina [42]. In [41], PRX-C contained residues 1036-1383 of rat L-PRX; in this particular region, a truncation has been linked to Charcot-Marie-Tooth disease type 4F, an incurable peripheral neuropathy.

SEC-SAXS data was collected for PRX-C (Figure 10A) in [42]. As expected, the AlphaFold prediction is mostly devoid of regular secondary structures and has a low confidence score (mean pLDDT < 50, Figure 10B). The structure does not fit the SAXS data and requires additional tools. The necessity for using ensemble-modeling methods for SAXS data is evident. ATSASfold provides the option to either select rigid domains that would not be modified, with flexible parts affected by ensemble-modeling, or to submit the whole chain as fully flexible to the program EOM. Using the second option for PRX-C, we obtained results in line with [41] (Figure10C-E).

## 6. Discussion

The examples presented above show both the strengths and the limitations of ATSASfold. One of its main purposes is to provide a continuous workflow between the initial analysis of experimental SAXS data, the comparison with an AF model and, if required, the selection and execution of an appropriate structure-refinement procedure.

Several existing programs and web servers address related problems, although differing in their main purposes. The most recent example is BilboMD [15]. This web-server provides an integrated workflow for conformational sampling using molecular dynamics, SAXS fitting with FoXS and the selection of multistate models with MultiFoXS [12]. It also provides tools for defining rigid and flexible regions, including their identification from the AlphaFold PAE score. Another recent development, SAXS-A-FOLD, starts from an AlphaFold or a user-provided structure and generates conformational ensembles by sampling selected flexible regions, followed by fitting of the resulting models against experimental SAXS data [13]. These developments illustrate a general trend towards combining predicted structures and SAXS modelling within integrated web-based workflows.

ATSASfold approaches this problem from a different direction. It is not dedicated to a particular refinement algorithm, but to the selection of an appropriate modelling strategy. A disagreement between an AlphaFold structure and a SAXS curve can have different structural origins and therefore does not always require the same treatment. Small changes within an otherwise compact structure can be approached using normal-mode refinement, whereas a change in the relative arrangement of rigid domains is better suited to rigid-body modelling. Conformational heterogeneity, by contrast, requires an ensemble description. ATSAS provides separate methods for these cases, including rigid-body modelling (BUNCH/CORAL/SASREF), SREFLEX normal-mode refinement and EOM ensemble modelling. The role of the ATSASfold classifier is to assist the user in selecting between these alternatives. In addition, ATSASfold provides the required input for the corresponding modelling procedure.

The four-class classifier (Direct fit, Rigid-body, Normal mode and Ensemble) is designed as a tool to support decisions rather than to provide a definitive structural classification. Indeed, the boundaries between these classes are not sharp. For example, a structure that requires only a small change in the relative position of two domains may have characteristics intermediate between the Direct Fit and Rigid-body classes. Similarly, the conformational changes that can be described by normal modes may overlap with systems displaying broader flexibility. For this reason, the classifier reports class probabilities in addition to the selected class. Cases in which two or more classes have similar probabilities should be considered ambiguous, and alternative refinement approaches may need to be tested. For this purpose, even with the Advanced mode turned off, the user can override the ATSASfold suggestion and proceed with the preferred modelling path.

A second limitation arises from the data used to train the classifier. The training data are a subset of the SASBDB, which is a curated but heterogeneous archive containing data collected at different facilities, at different times and using different experimental and processing procedures. The deposited entries also differ in sample quality and may lack some metadata. The structural modelling performed by the entry authors is evidently very different as well. Such differences may and do introduce biases into the classifier. The class labels, moreover, derive from the deposited models and from the automated refinement run described in the Appendix, rather than independently established ground truth; the training set therefore does not constitute an experimentally controlled benchmark. Further development would therefore benefit from a larger training set and from validation on independent datasets that were not involved in the classifier development.

In a more general context, a few routine remarks on biological SAXS data analysis should be made in the context of the ATSASfold workflow. The first concerns the limited structural information provided by SAXS. One should always remember that different three-dimensional structures can produce nearly identical scattering profiles, reflecting the inherent ambiguity of the inverse scattering problem [43]. Consequently, improved agreement with the experimental curve does not, by itself, demonstrate that the resulting model represents a unique solution structure. Increasing the number of adjustable structural parameters may also improve the fit without necessarily increasing the biological correctness of the model. To address this, ATSASfold specifically warns when the number of fitted parameters in rigid body modelling exceeds the estimated number of Shannon channels available in the data.

Likewise, the second comment concerns SAXS-AF fits. A poor agreement between an AlphaFold model and experimental SAXS data is not automatically evidence for a conformational difference. Disagreement may also come from the sample or experimental data. Aggregation, interparticle interactions, concentration effects, inaccurate buffer subtraction or radiation damage can affect SAXS profiles and consequently influence structural modelling. Differences may also result from an incorrect oligomeric state, missing residues, post-translational modifications, bound ligands or other components that are not represented in the input model. The primary analysis performed by ATSASfold helps to identify some of these problems before structural refinement is attempted. Yet, it cannot exclude all possible experimental or sample-related effects. The refinement recommendation should therefore always be considered together with information about sample preparation, experimental conditions and the biological and chemical properties of the studied protein (and buffer).

These limitations are well recognized in the guidelines for SAXS-based structural modelling, which discuss evaluation of model ambiguity and promote the use of additional experimental or biochemical information whenever possible [44]. The structures generated through ATSASfold-assisted fits should therefore be regarded as models consistent with the SAXS experiment rather than uniquely determined structures.

The present version of ATSASfold is restricted to protein monomers. This restriction simplifies the classification problem, but also excludes many SAXS applications involving oligomers, protein complexes and mixtures of different species. In such systems, disagreement with an AlphaFold model may be caused by changes in stoichiometry or association state rather than by conformational rearrangement of an individual chain. Extension to multimeric systems would therefore require additional analysis steps and routines. Other possible developments include expansion and independent validation of the classification dataset and an improved use of information provided by structure-prediction methods when defining domains and flexible regions.

## 7. Availability

The current version of ATSASfold web-server runs at EMBL Hamburg (https://www.embl-hamburg.de/biosaxs/atsasfold/). Registration with an email is required. In order to run ATSAS online projects, the emails on both services must match and be synced by pressing the respective button in ATSASfold. The AlphaFold runs are possible only in the local EMBL network, e.g. at the P12 beamline. The server and ATSAS online use ATSAS version 3.2.2 [4].

The manual segmentation tools’ source is openly available for download via Git or Zenodo [36]. Also, the code for ML classifier model, together with the model weights, and the description of the latest dataset, is available at [45].

Users are invited to submit bug reports & suggestions using online form at the website or directly by email to.

The SASBDB codes for the examples used in this manuscript were SASDFP8 (carbonic anhydrase II), SASDP39 (ATE1) and SASD2N7 (PRX-C), and are openly available at the SASBDB website (https://www.sasbdb.org).

## 8. Conclusions

The ATSASfold web-server provides the workflow for connecting experimental SAXS data with AlphaFold models and ATSAS refinement tools. It bridges the gap between SAXS data interpretation and AlphaFold-based structure prediction, offering a user-friendly, guided workflow for structural biologists, for whom deep expertise in SAXS or computational modelling are not required. The server performs primary SAXS analysis, evaluates the agreement between predicted structures and solution scattering data, and suggests suitable modelling strategies when discrepancies are detected. Its ML-driven classifier provides recommendations for structure refinement, with an aim to reduce the trial-and-error process. By integrating ATSAS, AlphaFold, and domain-specific tools, ATSASfold simplifies complex workflows keeping the access to advanced refinement strategies. Still, users should validate the ML-guided decisions, particularly in cases with low classification confidence. As the tool evolves alongside AlphaFold3 [7] and new SAXS methodologies, future iterations could further reduce required user input during data analysis.

An advantage of ATSASfold is its modularity: users can switch between its “guided” and “advanced” modes, allowing flexibility for both novices and experts. The inclusion of visualization tools (e.g., Kratky/PDDF plots, PAE/pLDDT-based domain segmentation) helps interpretability, making it easier to assess data quality and model agreement with experimental observations. The presented examples show that ATSASfold can simplify SAXS interpretation for new or inexperienced users while preserving access to established ATSAS methods. In this way, ATSASfold may help SAXS users move more efficiently from beamline measurements to structural hypotheses.

## Appendix A. ML-classifier architecture

In this section additional details are given for the ML-classifier that assists user decision at the refinement stage. As described in the main part, its task is to assign the project to one of four classes – Direct fit, Rigid-body, Normal-mode, or Ensemble basing on the various features extracted from SAXS experimental data, AlphaFold predicted structure and their agreement. For this purpose an XGBoost classifier [33] was trained on the dataset composed of processed SASBDB depositions augmented with a few additional inputs.

### A1. Dataset and features engineering

The main source of the training data was SASBDB - Small Angle Scattering Biological DataBank, a curated repository of freely accessible and fully searchable SAS experimental data. SAS curves are deposited together with the relevant experimental conditions, sample details, instrument characteristics and derived models. The quality of deposited experimental data and the accuracy of models obtained from SAS and complementary techniques is assessed by the site developers [17] (EMBL Hamburg). SASBDB exposes REST API to conveniently download all public data in JSON/XML formats. This databank has recently bypassed 5000 depositions [46], and as of November 2026, it contains more than 5700 entries with solution SAXS/SANS data of protein monomers, complexes, RNA and some nanoparticles.

To produce the training dataset for ATSASfold classifier, we have downloaded all the protein monomer entries from SASBDB, obtaining roughly 1500 sets. Each entry consists of a SAXS curve, AA sequence of the protein, reported PDDF, *R_g_* and *D_max_*.

A first-round assigning of classes to the entries was the following. If the entry depositor reported the use of SREFLEX, SASREF, BUNCH/CORAL, EOM or other tools like MultiFoXS or AMBER, etc, together with an improved *χ*^2^, the entry was attributed to the corresponding class. In case *χ*^2^ agreement was not acceptable, the class was not assigned. Also, models tagged as IDPs by SASBDB tagging system were assigned directly to Ensemble class.

The main analysis and class assignment pipeline consisted of several steps, some of which were automated. First, for each entry a respective AF model together with PAE and pLDDT scores was obtained by running inference at the EMBL HPC cluster [47]. Next, the fits quality of the atomic structure to the experimental curve was evaluated (via CRYSOL runs at the Hyde cluster in EMBL Hamburg). If 2 out of 3 CRYSOL fit quality evaluation scores (*χ*^2^, CorMap, Anderson-Darling) were in the green zone, the entry was automatically assigned to Direct fit class. Further, quality of the fit was additionally evaluated by using CRYSOL scores together with 3 other metrics (*D_log_*, *V_r_* and *D_Rg_*, see below). If, again, two of them were acceptable, the Direct fit class was also assigned.

The remaining entries were distributed between the other three classes. In order to do this, SREFLEX runs on the cluster were submitted and evaluated using an automated pipeline. In case of a very good obtained fit quality, *χ*^2^<1.1, the Normal-mode class was assigned. In parallel, the Chainsaw and Merizo methods were used to determine protein domains and linkers, and the results were also submitted as CORAL runs. Again, in case of a good fit quality, the Rigid-body class was automatically assigned. Between Rigid-body and Normal-mode the priority was given to later one. The scheme of this approach is thus summarized on Figure 11.

**Figure 11.**
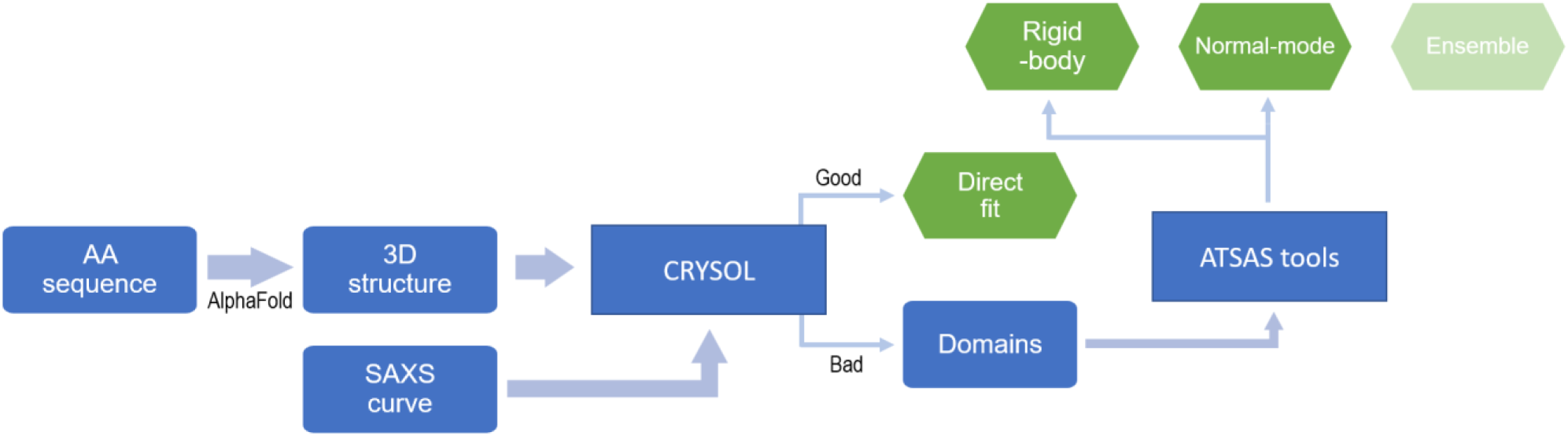
Scheme of choosing a category for the entry for the cases when it was not classified according to SASBDB entry data.

After this procedure, approximately 750 entries with selected class labels were obtained. As it turned out during training, a considerable number of misclassifications still existed and required manual attention. In order to do this, we have analysed several lists of entries, ordered by fit quality and other features. This allowed to identify obvious (during manual review) errors in classifications, and also analyse some outliers in detail. For most of these cases several different ATSAS online runs were manually performed in order to clarify which method improves the fit. Additionally, some cases required analysis of the SASBDB-reported paper in search for hints on possible class selection. We expect to have updates to the respective repository with new datasets as some manual processing continues. Furthermore, as additional entries are added to SASBDB, the training dataset will be expanded.

Next, the feature generation pipeline has been developed. It involves running different deterministic scripts on the data. Up to this moment, apart from AF structure and scores, each entry had a CRYSOL fit, and regularly also the CORAL/SREFLEX run data. The pipeline consists of further steps:

1. Extracting PAE and pLDDT scores and some of their characteristics (e.g. mean/min/max pLDDT, PAE, and more complex features).
2. Extracting AA sequence embedding vector after ProtT5 or ESM2 inference [48,49].
3. Principal components analysis (PCA) classification of embeddings. This classification was performed in supervised mode (labels-aware), and applied to reduce the embeddings feature size to 64 floating point values.
4. Obtaining 6 different fit quality metrics from the CRYSOL fit, some of them not depending on experimental errors reported.
5. Estimating *R_g_*, *D_max_*, *I*(0) values by running autorg and datgnom tools. Thus, also a separate PDDF function was obtained and saved.
6. Obtaining and saving the dimensionless Kratky plot.
7. Extracting SAXS/PDDF/Kratky curves features.
8. Extracting domain/linker related features.

The obtained features were grouped into a feature set and used further for training. Overall, 4 broad categories can be distinguished. These categories describe sequence information, confidence in the predicted structure, experimental SAXS and model–experiment agreement, and domain organization. Some categories share information sources, particularly those comparing the AF3 model with experimental SAXS data. The feature groups are summarized next. While describing specifically each feature is avoided here, we refer the reader to [45] for more details.

#### AA sequence features (64 features)

Sequence information was represented using protein language model embeddings averaged over residue positions. ProtT5 and ESM-2 were evaluated: ProtT5 gave a small accuracy improvement, while running ESM-2 required fewer computational resources. The current training feature set uses an ESM-2 embedding reduced to 64 components via PCA; 16, 32 and 128 components were also tested.

#### AF3 confidence features (18 features)

This group represents local structural confidence and uncertainty in relative residue placement. It includes the AF3-reported disordered fraction, mean residue pLDDT, and mean pLDDT in eight successive sequence segments. PAE-based features show intrachain uncertainty across different separation ranges, together with ratios and differences comparing local and longer-range relationships. Thus, these features provide information on the distribution of confidences along the proteins sequence and uncertainties in relationships between the distant residues.

#### SAXS and model–experiment agreement features (65 features)

This group contains the summarized experimental scattering data and also reflects its agreement with the predicted structure. It includes experimental estimates of *R_g_*, *I*(0) and *D_max_*, corresponding features of the AF model, and ratios quantifying size disagreement. The pair-distance distribution is represented by nine normalized area bins. Dimensionless Kratky descriptors summarize its binned profile, data coverage, peak properties and high-*q* behavior. These features may indicate differences in particle shape or flexibility. The agreement between experimental and modeled scattering is described by CRYSOL *χ*^2^ scores and uncertainty-normalized residual summaries across different *qR_g_* ranges. Additional features characterize experimental signal-to-noise, reported parameter uncertainties and the starting points selected for Guinier fitting and SAXS processing.

#### Domains organization features (16 features)

This group describes the Chainsaw-derived domain organization through domain counts, including the number and sizes of domains and the number and lengths of linker regions between domains or the ends of the sequence. Spatial relationships between domains are summarized by centroid distances and contact densities calculated from C_α_ coordinates. Within-domain and between-domain PAE summaries characterize confidence in individual domains and their relative placement. Together, these descriptors reveal domain organization, packing and uncertainty in interdomain arrangement.

### A2. Additional fit quality metrics

As mentioned above, additional fit quality metrics were calculated for each entry. The idea behind their implementation was to evaluate how well does the fit follow the experimental curve, without using reported experimental errors. It is worth noting once again, that being an open repository, SASBDB does not introduce strict rules or control on deposited data, thus some entries may report overestimated or underestimated experimental errors. This then affects the classifiers’ performance on CRYSOL *χ*^2^ scores used as features.

Thus, three supplementary metrics per curve were calculated: *D_log_*, *V_r_* and *D_Rg_*. The *V_r_* metric is the volatility of ratio metric defined in [50] and adapted for the pipeline here. The *D_log_* and *D_Rg_,* as well, are pipeline-defined fit quality features, with *D_Rg_* being merely the absolute value of the logarithm of ratio of gyration radii calculated from experimental and fit curves. The *D_log_* feature, on the other hand, is the numerical integral of *d*(*q*) function, where

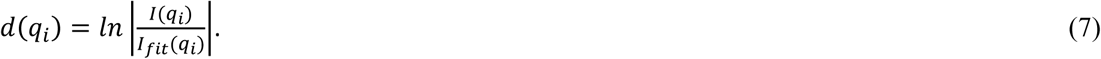

### A3. Data augmentation

Additionally, we have added simulated data to the dataset basing on PDB entries used to test and demonstrate SREFLEX method performance [51]. Overall, 44 PDB codes pairs were downloaded. The extracted AA sequences were used to generate the AlphaFold models. Out of each pair we have then taken the structures that differed more by RMSD from the AF models and generated SAXS curves using CRYSOL, with background and noise added. Entries for which CORAL refinement resulted in a substantial improvement in fit quality were assigned to the Rigid-body class. Overall, 44 more non-Direct fit, non-Ensemble entries were considered. Out of them, only 6 entries were added to final training set.

These simulated data entries were restricted only to training and were not included into evaluation of model performance.

### A4. Feature selection and resulting dataset

As the result, our dataset consisted of 738 entries, of which 6 were simulated data. The Direct fit and Ensemble classes form the majority of the dataset (200 and 316 entries, correspondingly), while Normal-mode and Rigid-body are the minor classes (145+4 and 71+2 entries).

For the feature selection we have run a few tests, including exploratory comparison of datasets containing just “single source” features: ESM-2 processed embeddings dataset (64-float vector), SAXS features dataset (61 features), AF-inferred structure dataset (38 feature) and their combinations, including one with all three combined. While doing investigations on possible features to extract for each entry, more than 270 separate features were derived. During training the features having more impact on classification quality were selected and used further.

The features used during training are summarized as feature groups in Table 1. The datasets described were used for training the ATSASfold ML classifier.

**Table 1.** List of feature groups used for training the ATSASfold ML classifier. The first three columns display sources of data for the features and their intersections.

| AF | SAXS | Domains | **Feature group** | **Number of features** |
|---|---|---|---|---|
|  |  |  | ESM sequence representation | 64 |
|  |  |  | AF3 disorder and residue confidence | 10 |
|  |  |  | PAE by sequence separation | 8 |
|  |  |  | Experimental and model size | 8 |
|  |  |  | Pair-distance distribution profile | 9 |
|  |  |  | Dimensionless Kratky profile and coverage | 17 |
|  |  |  | Kratky peak and tail descriptors | 9 |
|  |  |  | CRYSOL residual profiles | 14 |
|  |  |  | Overall CRYSOL fit | 2 |
|  |  |  | Experimental quality and analysis window | 6 |
|  |  |  | Domain organization, contacts and linkers | 10 |
|  |  |  | Within-domain and between-domain PAE | 6 |
|  |  |  | **Total** | **163** |

### A5. Training XGBoost classifier

XGBoost belongs to a family of gradient boosting decision trees (GBDT) models. It iteratively fits decision-trees to the residuals of the previous models, thus continuously improving ensembles performance. During training, we have used stratified cross-validation to preserve class balance across folds. The tuning of hyperparameters (model selection) was carried out with Optuna [52], optimizing the balanced accuracy to reduce bias toward majority classes (Fixed, EOM). We also inspected overall accuracy, per-class precision and recall, and confusion matrices to asses where misclassifications occurred (as described above).

After selecting the best hyperparameters, the final model was refitted on all training-eligible data and exported together with metadata, metrics and, importantly, preprocessing steps to be reproduced by ATSASfold. Reproducibility checks were generated to verify successful migration.

### A6. Training results

Performance of the classifier was assessed using balanced accuracy score. Given the number of true positives (TP), true negatives (TN), false positives (FP) and false negatives (FN) per each class, one can calculate several metrics, with the balanced accuracy being the macro-average of recall scores per class. Thus, the expression for the metric used for training and hyperparameters optimization is derived from per-class recall:

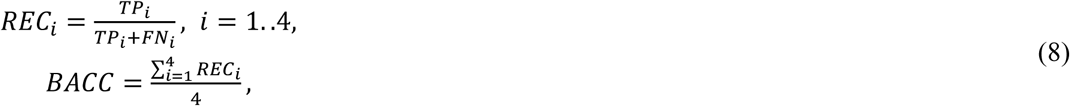

basically reflecting the probability of correct classification balanced over classes. Balancing is helpful for considered dataset, since Rigid-body and especially Normal-mode classes are the minority classes. The plain accuracy score, also reported, is regularly 0.05-0.1 higher after training. Additional scores reported are the per-class metrics (precision, recall and F1 score), and also macro and weighted metrics, confusion matrix.

As stated above, it was interesting to compare the performance of ablated datasets, containing grouped-feature only, and their combinations. The resulting scores are reported in the Table 2.

**Table 2.** Comparison of classifier’s performance scores in the full mode and with ablations of different feature groups.

| **Model name** | **Accuracy** | **Balanced ACC** | **Macro-avg precision** | **Macro-avg F1** | **MCC** |
|---|---|---|---|---|---|
| ESM | 0.6940 | 0.6309 | 0.6668 | 0.6447 | 0.5509 |
| AF | 0.7473 | 0.6733 | 0.7010 | 0.6827 | 0.6323 |
| SAXS | 0.7104 | 0.6263 | 0.6680 | 0.6382 | 0.5741 |
| AF+SAXS | 0.7842 | 0.7082 | 0.7358 | 0.7168 | 0.6855 |
| ESM+AF | 0.7473 | 0.6820 | 0.7100 | 0.6932 | 0.6305 |
| ESM+SAXS | 0.7773 | 0.7095 | 0.7552 | 0.7261 | 0.6735 |
| ESM+AF+SAXS | 0.7951 | 0.7281 | 0.7612 | 0.7414 | 0.6998 |

## Notes

### Competing Interest Statement

The authors have declared no competing interest.

